# Modulating the selective utilization of carbon sources by engineering the 3^rd^ and 4^th^ helices of the DNA-binding domain of catabolite control protein A (CcpA) in *Bacillus licheniformis*

**DOI:** 10.1101/2021.05.17.444589

**Authors:** Yupeng Zhang, Youran Li, Fengxu Xiao, Hanrong Wang, Liang Zhang, Zhongyang Ding, Zhenghua Gu, Sha Xu, Guiyang Shi

## Abstract

The gram-positive bacterium *Bacillus licheniformis* exhibits obvious selective utilization on carbon sources. This process is mainly governed by the global regulator catabolite control protein A (CcpA), which can recognize and bind to multiple target genes widely distributed in metabolic pathways. Although the DNA-binding domain of CcpA has been predicted, the influence of key amino acids on target gene recognition and binding remains elusive. In this study, the impact of Lys31, Ile42 and Leu56 on in vitro protein-DNA interactions and in vivo carbon source selective utilization was investigated. The results showed that alanine substitution of Lys31 and Ile42, located within the 3^rd^ helices of the DNA-binding domain, significantly weakened the binding strength between CcpA and target genes. These mutations also lead to alleviated repression of xylose utilization in the presence of glucose. On the other hand, the Leu56Arg mutant in the 4^th^ helices exhibited enhanced binding affinity compared with that of the wild-type one. When this mutant was used to replace the native one in *B. licheniformis* cells, the selective utilization of glucose over xylose increased. The above research results are helpful for a deep understanding of how microorganisms can flexibly sense and adapt to changes in the external environment. Additionally, they can provide important theoretical basis for the rational design of biomass utilization and environmental adaptability of *B. licheniformis* cell factories.

**Importance:** *Bacillus licheniformis* is widely used in producing various valuable products, such as α enzymes, industrial chemicals and biocides. The carbon catabolite regulation process in the utilization of raw materials is crucial to maximizing the efficiency of this microbial cell factory. CcpA plays an important role in this process. This study represents a new paradigm to investigate the structure–function relationship in CcpA by fluorescence polarization experiments in vitro. The results also uncover key amino acids in the DNA-binding domain that affect the selective utilization of carbon sources. These results provide a theoretical basis for the rational design of industrial microorganisms.

## Introduction

Microorganisms have evolved a myriad of strategies to adapt to complex environments. In firmicutes, the regulation of carbon resource utilization is mainly governed by the global regulator catabolite control protein A (CcpA), a LacI-GalR family protein (1). As this regulator can have direct or indirect effects on multiple genes involved in both catabolism and anabolism, its specific function and mechanisms of action have attracted increasing attention. The domains of CcpA in *Bacillus subtilis* and *Bacillus megaterium* have been characterized (2, 3). Researchers have suggested that bacterial CcpA contains two domains: a DNA-binding domain and a core domain (4, 5). The DNA-binding domain is mainly responsible for the recognition of nucleic acids, and the core domain is mainly responsible for cofactor HPr (histidine-containing protein) or Crh (Carbon flux regulating HPr) binding (6, 7). With the increasing availability of “multi-omics” information, CcpA’s binding site was found to be widely distributed (8). However, the relationship between the structure and function of CcpA, especially the influence of key amino acids on target gene recognition and binding, is still elusive. On this basis, the carbon metabolism regulatory networks centered on CcpA are only fully comprehended by considering its structural context.

The current understanding of the regulatory function of CcpA in gram-positive bacteria is mainly based on research conducted in *Bacillus subtilis* and *Bacillus megaterium*, in which the function of CcpA has been appraisal by examining effects of amino acid substitutions (2, 3, 6). In *B*. *megaterium*, the mutation of some amino acids was used to research the regulatory effect of CcpA on growth and catabolite repression. Mutations of Glu77, Ile227, Asp275, Met282 and Thr306 show glucose-independent regulation (2). In both studies, the chosen amino acids are mainly located in the core domain and are highly conserved.

*B. licheniformis*, a gram-positive bacterium with great application potential, is not only used for a wide range of applications in the field of fermentation but also an excellent platform for exogenous gene expression (9, 10). In the field of fermentation, because of the unique advantages of *B. licheniformis* (a moderate growth rate and sufficient protein folding activity (11)), *B. licheniformis* is used to produce bacitracin (12), poly-γ-glutamic acid (13), amylase (14), and alkaline protease (15), among others. However, in industrial fermentation, the presence of a preferred carbon source, such as glucose, inhibits the utilization of a nonpreferred carbon source until the preferred carbon source has been exhausted (16). This phenomenon is called carbon catabolite repression (CCR), of which glucose-lactose diauxie in *Escherichia coli* is a classic example (17). CCR, one of the most widespread mechanisms by which microbes adapt to a changing environment, takes advantage of protein synthesis (18). Generally, the major determining factor in the microbial growth rate is the selection of a preferred carbon source (17). In addition, the presence of a preferred carbon source will also cause other metabolic changes beyond CCR (19).

Xylose can be utilized by *B. licheniformis* (20). and is a readily available, abundant, inexpensive, and renewable resource of biomass (21, 22). In the hydrolytic products of lignocellulosic biomass, xylose is not only mainly contained, but glucose is also one of the main hydrolytic products of lignocellulosic biomass (23). The utilization of xylose is repressed because of the existence of glucose under the regulation of CcpA (24). Therefore, the adjustment of regulation in CcpA may affect the preference of xylose.

Here, we show that some amino acid mutations in the DNA-binding domain can influence the binding ability of CcpA, thus causing changes in its regulatory function in *B. licheniformis*. The utilization of xylose was repressed by the binding of CcpA with nucleic acid at its binding sites. The effect of amino acid mutations in the DNA-binding domain of CcpA on xylose utilization was investigated. These findings may help us understand how CcpA regulates the utilization of xylose in *B. licheniformis* and open new possibilities for the collaborative utilization of the preferred carbon source and xylose.

## Results

### Homology modeling of *B. licheniformis* CcpA and sequence alignment analysis

Diversified cre sites that can bind CcpA and changes in these sites can effectively influence the binding ability of CcpA. These changes influence the regulation of metabolism by CcpA (25, 26). To explore whether changes in the amino acids of the CcpA protein would also affect the function of CcpA, first, a homology model of CcpA was built by SWISS-MODEL (Fig. 1A). The obtained model of CcpA in *B. licheniformis* showed high homology (87.01%) with the model (1ZVV) we used. Verify3D showed that at least 80% of the amino acids were scored (27). In addition, ERRAT showed that the quality factor was 95.5 (28). These results indicated that the modeled structures of CcpA can be used for subsequent analyses. The model shows that the DNA-binding domain of *B. licheniformis* CcpA contains four α-helices. In previous studies, Arg22 and Leu56 at the 2^nd^ and 4^th^ α-helix had been demonstrated to play an important role in interaction of CcpA with cre sites (29). To further confirm the key amino acids that can influence the regulation of CcpA, we analyzed CcpA from seven closely related *Bacillus* species. Identical and similar amino acids in the DNA-binding domain are marked in blue, and conserved amino acids are marked in red (Fig. 1B).

**FIG 1.**
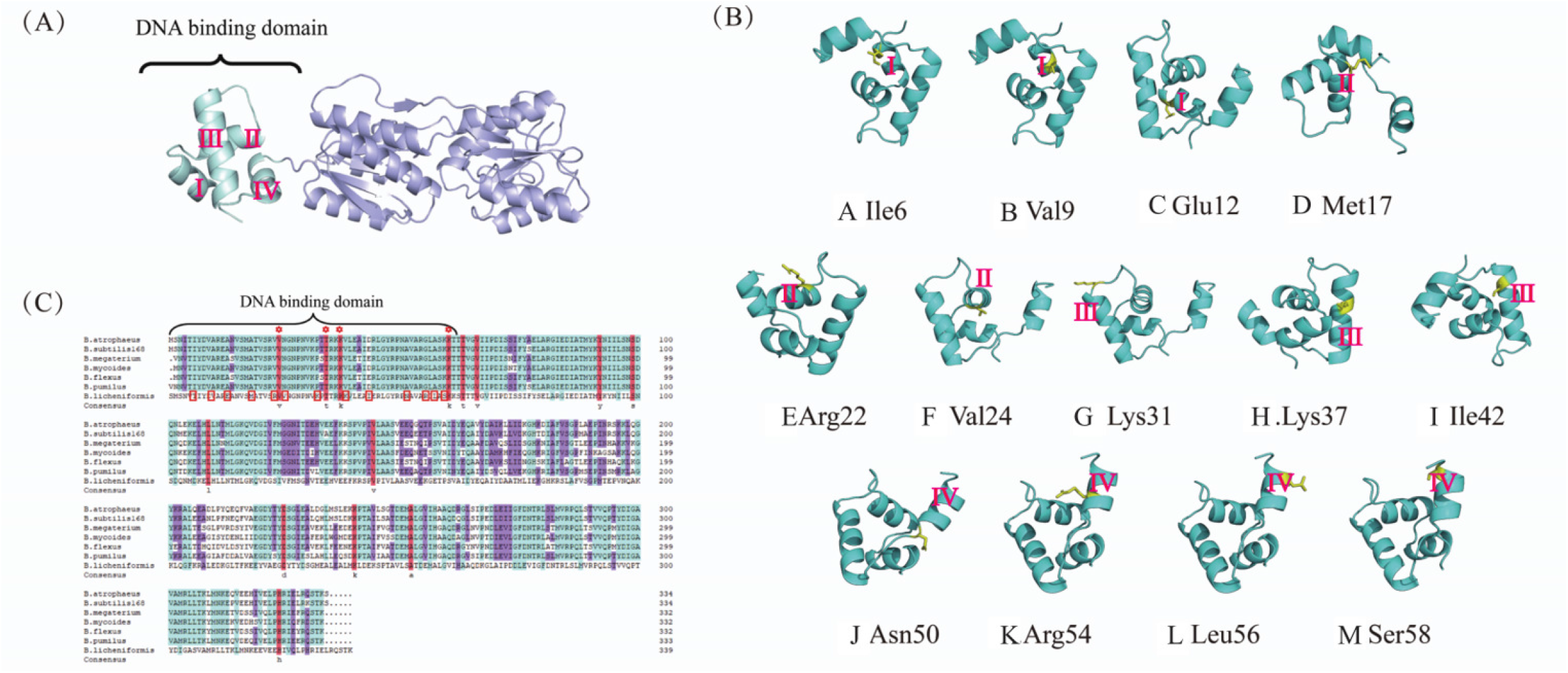
Selection of the key amino acids in CcpA DNA-binding domain. (A) The structure of the whole CcpA was divided into two domains, DNA-binding domain (marked by blue) and cofactor-binding domain (marked by purple). (B) The location of amino acids we selected in DNA-binding domain. The amino acids we selected and side chain were marked by yellow. (C) The location of amino acids in others *Bacillus*. The conserved amino acid in different strains were marked by red. The amino acids we selected were marked by red box. The accession numbers used for acid sequence analysis were WP_010789522.1 (B. atrophaeus), WP_025909479.1 (B. flexus), WP_014458003.1 (B. megaterium), EEK71284.1 (B. mycoides), WP_061409141.1 (B. pumilus) and WP_003229285.1 (B. subtilis168).

The structures of CcpA subunits are flexible, and the structure of CcpA changes to fit nucleic acids for binding. The main change in CcpA when it binds nucleic acids is a change in the angle between different α-helices of the DNA-binding domain (29). In *B. subtilis*, the mutations of Asn49 and Asn50, which are next to the hinge helix, were likely to directly or indirectly affect DNA-binding (30). Interestingly, these amino acids were located at one end of the 4^th^ helices. The key amino acids that cause this change in the angle between different α-helices may be located at both ends of the α-helix. Alignment of the amino acid sequences of *Bacillus*, Val23, Thr33, Lys36 and Lys59 were strictly conserved in the DNA-binding domain. These amino acids were located at the end of helices and the midpiece of helices. They were believed to be the essential residues that are important for the stability and activity of CcpA (29). Hence, the amino acid in DNA-binding domain plays an import role in recognition and binding. On the other hand, the amino acids near conservation may affect the function of CcpA. Based on the above information, the amino acids Ile6, Va9, Glu12, Met17, Arg22, Val24, Lys31, Lys37, Ile42, Asn50, Arg54, Leu56, and Ser58 were chosen as target positions, which are mainly located at either end of the α-helix or near the conserved region, for further examination, as shown in Fig. 1C.

### Site-directed mutagenesis, clone expression and purification of the *B. licheniformis* CcpA protein

Based on the above analysis, the putative amino acids within the α-helix were replaced by site-directed mutagenesis. In a previous study, an alanine scanning peptide library was used to estimate the specific amino acids that were correlated with stability and function (31). To test the function of amino acids, some residues were replaced by alanine using site-directed mutagenesis. Schumacher MA confirmed that Leu56 identifies the conserved domain of cre during binding with nucleic acids(29). The characteristics of Leu were opposite those of Arg. Leu56 was mutated to Arg. The sequencing results showed that all the mutations were in agreement with the sequences shown in Table 2. Therefore, we obtained 13 CcpA mutants. All mutant proteins were expressed and purified in host *BL21* bacteria using the pET28a vector, and 35-67 mg of each purified protein was obtained. The mutant proteins migrated at their expected size as judged by SDS-PAGE (Fig. 2).

**FIG 2.**
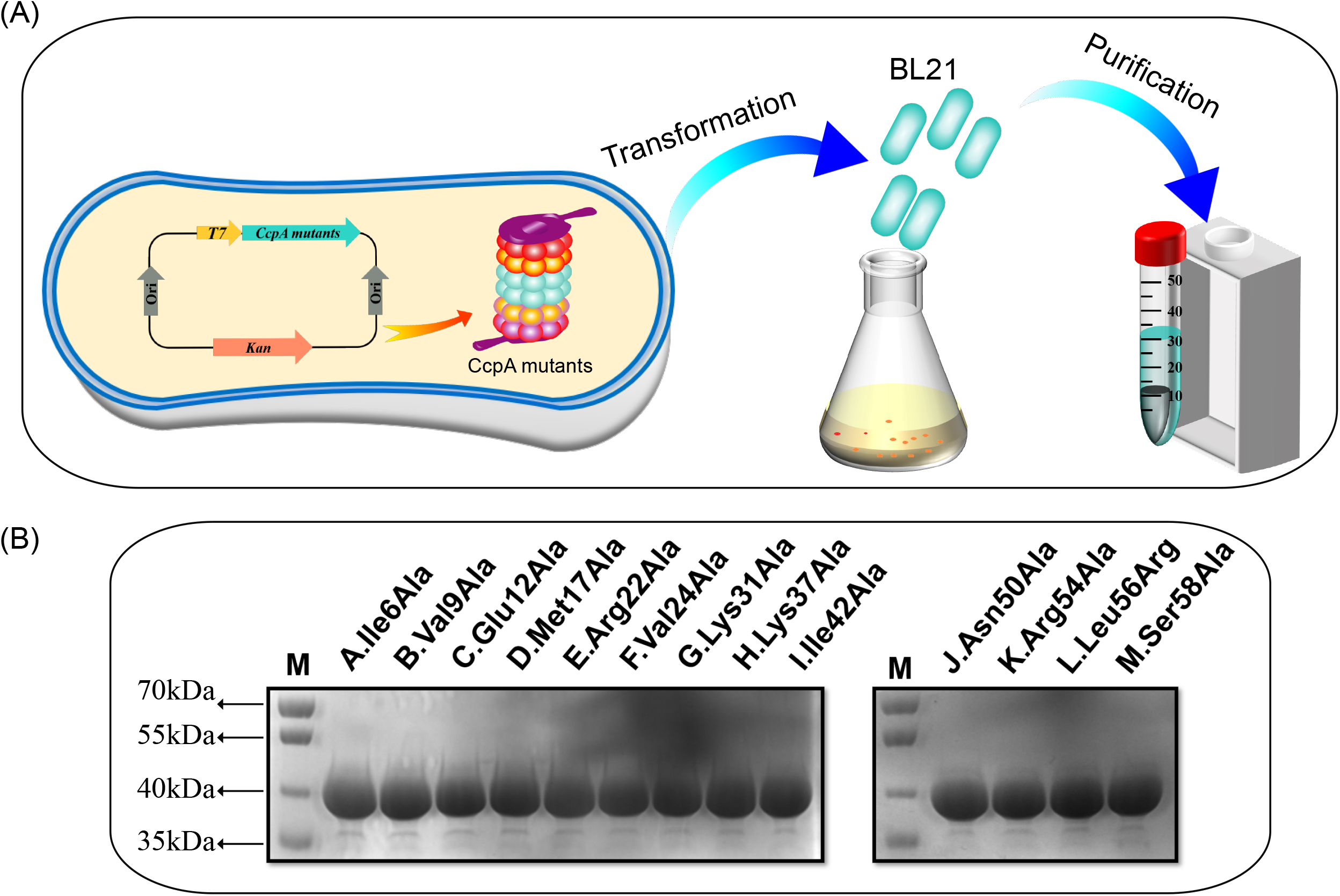
The mutation and purification of CcpA. (A) Schematic of heterologous expression and purification for CcpA mutants. (B) The purification of CcpA mutants. CcpA mutants were purified and then detected by SDS-PAGE.

### The binding ability of CcpA to cre sites changed due to the amino acid mutation

Previous studies had demonstrated that CcpA can directly and indirectly regulate the transcription of several essential genes by binding to *cre* sites (32). Therefore, the ability of CcpA mutants to bind cre sites was first evaluated in vitro. The proteins and the DNAs were mixed and subjected to fluorescence polarization (FPIA) and electrophoretic mobility shift assays (EMSA), the results of which are shown in Fig. 3. A bar chart exhibited fluorescence polarization after binding of the mutant proteins compared to the wild-type protein to the cre site (Fig. 3A). It is clear from the experimental results that the fluorescence polarization of Lys31Ala, Ile42Ala and Leu56Arg were significantly shifted compared with that of other mutants. Leu56Arg binds with higher affinity to the cre site than the negative control (wild-type CcpA). In contrast, Lys31Ala and Ile42Ala bind with lower affinity to the cre site than the negative control. To verify the above results, we examined the binding of CcpA mutants to the cre site by using electrophoretic mobility shift assays (EMSAs) (Fig. 3B). As expected, a substantial DNA-binding shift was observed in the Leu56Arg compared with the negative control. However, no obvious DNA-binding shift was detected for Lys31Ala and Ile42Ala. Together, these results verified that the amino acids that we chose in the DNA-binding domain play an important role in cre recognition.

**FIG 3.**
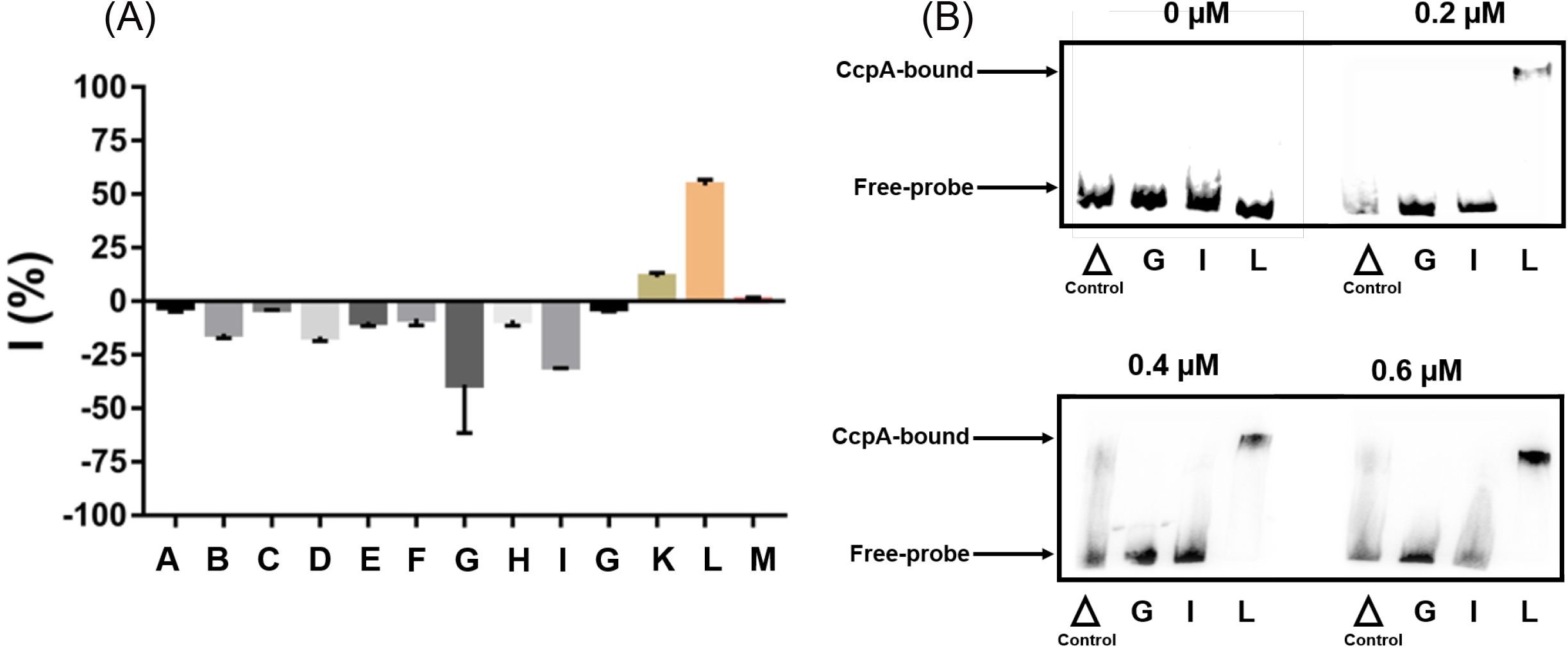
Evaluation of nucleic acids biding ability between different CcpA mutants. All experiments were performed in triplicate. (A) Screening the CcpA mutants by FPIA. “I” is defined to describe the difference in polarization value between the mutant and the wild-type CcpA. The value I is the polarization value of CcpA mutants minus wild-type CcpA. (B) EMSAs of wild type CcpA and CcpA mutants which we selected in (A) binding to the probe containing canonical cre site. The final concentration of 5’ labeled DNA probe used was 30 ng, and 0 to 0.6nm CcpA mutants were used. Wild-type protein was included as control. ***——P<0.001, **—P<0.01, *——P<0.05.

### The effect of CcpA mutants on glucose/xylose selective utilization

We first constructed a CcpA deletion mutant to investigate the effect of this protein on carbon source utilization and to provide a clear background for engineered CcpA proteins. This work was confirmed by diagnostic PCR and subsequent sequencing. The schematic of CcpA knockout strategy was shown in Fig. 4A. The *ccpA* deletion cassette exhibited a 2384 bp fragment in nucleic acid electrophoresis, as shown in Fig. 4B. The relative expression intensity of the *ccpA* gene was also documented at different times (Fig. 4C). Quantitative RT-PCR analyses of the levels of *ccpA* in the cells cultured for 4 h, 8 h and 12 h were performed. The *ccpA* expression level in the *ccpA* deletion strain was 1.6%, 0.4% and 4.4% of that in the wild-type strain. Compared with the original strain, the expression of origin ccpA was disrupted by exogenous gene successful.

**FIG 4.**
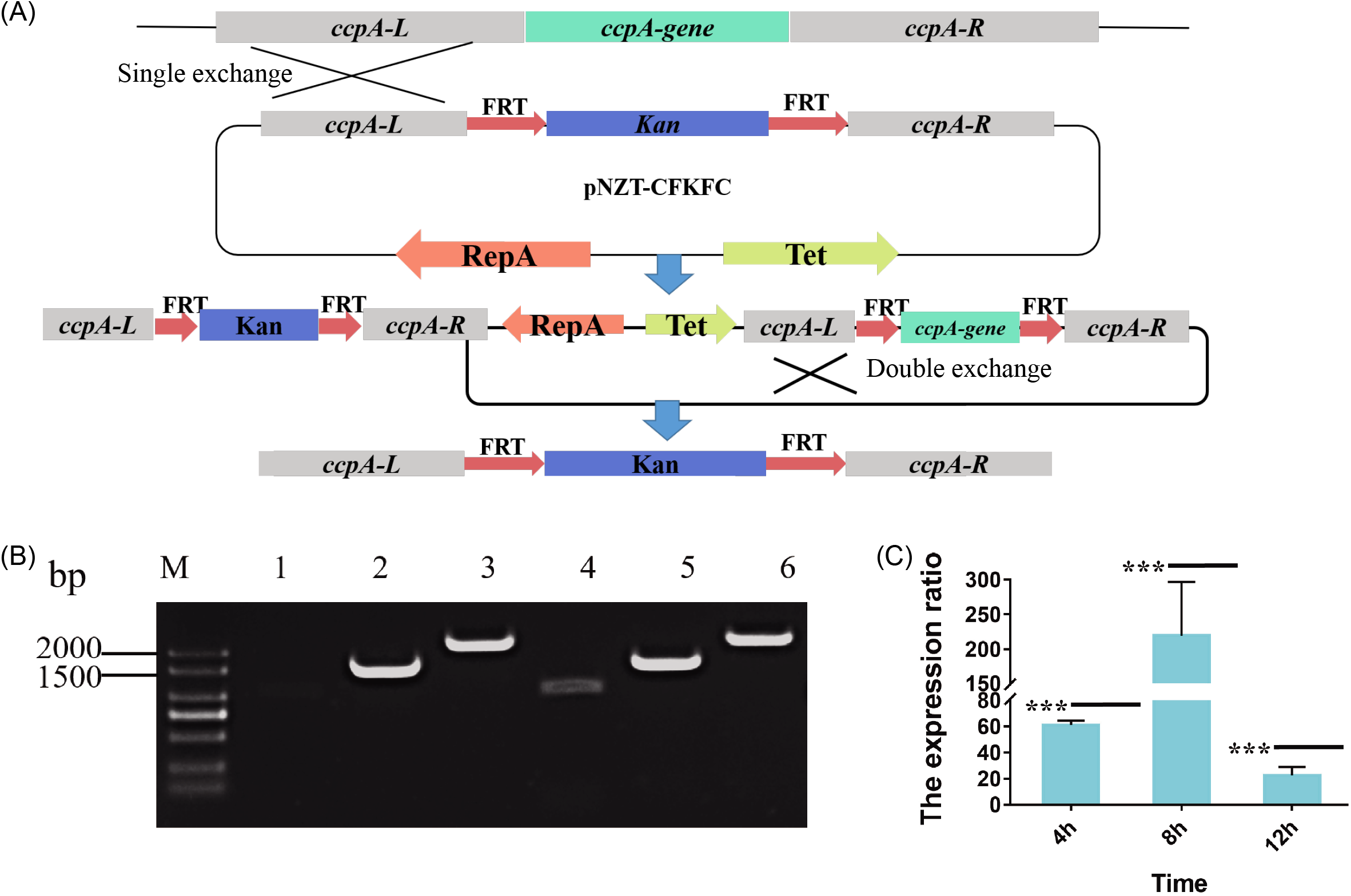
Verification of *ccpA* gene knockout. (A) The schematic of CcpA knockout strategy. *ccpA-L* and *ccpA-R* were left and right homology arms. The origin *ccpA* was replaced by *Kan* during homeologous recombination. (B) PCR products were used to verify the *ccpA* was replaced by foreign gene. The edited *ccpA* gene will have three bands (Lanes 4, 5, 6(1283bp, 1825bp and 2384bp)). Unsuccessful editing *ccpA* gene have two bands (Lanes 1, 2, 3). The sizes of the DNA markers are labeled on the left. (C) The expression ratio of *ccpA*. The expression ratio is the expression of wild-type *ccpA* divide by the expression of CcpA deletion. ***——P<0.001, **—P<0.01, *——P<0.05.

The repression of xylose utilization by glucose has been demonstrated in Bacillus strains, as shown in Fig. 5. Generally, CcpA binds cre sites in the presence of glucose, after which the expression of a xylose utilization gene is repressed (6, 20, 33). In the above studies, we demonstrated that the ability of a CcpA mutant to bind cre sites differed from that of wild-type CcpA (Fig. 5D). The mutant genes of *ccpA* were then expressed in *B. licheniformis CA* to explore their influence on carbon source selective utilization. Recombinant *bacteria* were cultured in TB medium suppled with xylose and glucose, the xylose consumption rate was used as an index. As expected, CcpA mutants, with diversified affinities to cre sites, exhibited significant differences in selective utilization between glucose and xylose (Fig. 6). The average specific consumption rate of xylose in the control (Wild-type *ccpA* was expressed in *B. licheniformis CA)* was approximately 0.25±0.1 g/ (L.OD_600_) in the presence of glucose. The results showed that the average specific rate of xylose consumption in the presence of the preferred carbon source was obviously higher in Lys31Ala and Ile42Ala CcpA than in the control group in the presence of glucose. As shown in Fig. 6E, the average specific consumption rates of xylose Lys31Ala in and Ile42Ala are approximately 1.5±0.3 g/ (L.OD_600_) and 1.2±0.02 g/ (L.OD_600_) at 21 h, respectively. The xylose consumption rate for Lys31Ala in and Ile42Ala increased by 5- and 3.8-fold relative to that for wild-type. In contrast, the average specific rate of xylose consumption of the Leu56Arg mutant was obviously decreased. The average specific consumption rate of xylose is approximately 0.04±0.003 g/ (L.OD_600_) in the presence of glucose. The xylose consumption rate for Leu56Arg decreased by 5.25-fold. Moreover, the ratio of glucose to xylose utilization was obviously different in strains expressing mutant CcpA. The ratio of glucose to xylose utilization was approximately 15 in the negative control group but approximately 4-fold lower in the strains expressing Lys31Ala and Ile42Ala CcpA. The ratio of glucose to xylose utilization was over 30-fold higher than that of the negative control in the strain expressing Leu56Arg CcpA. Furthermore, the biomass of strains that overexpressed the CcpA mutants was different from that of the strain expressing wild-type CcpA (Fig. 6D).

**FIG 5.**
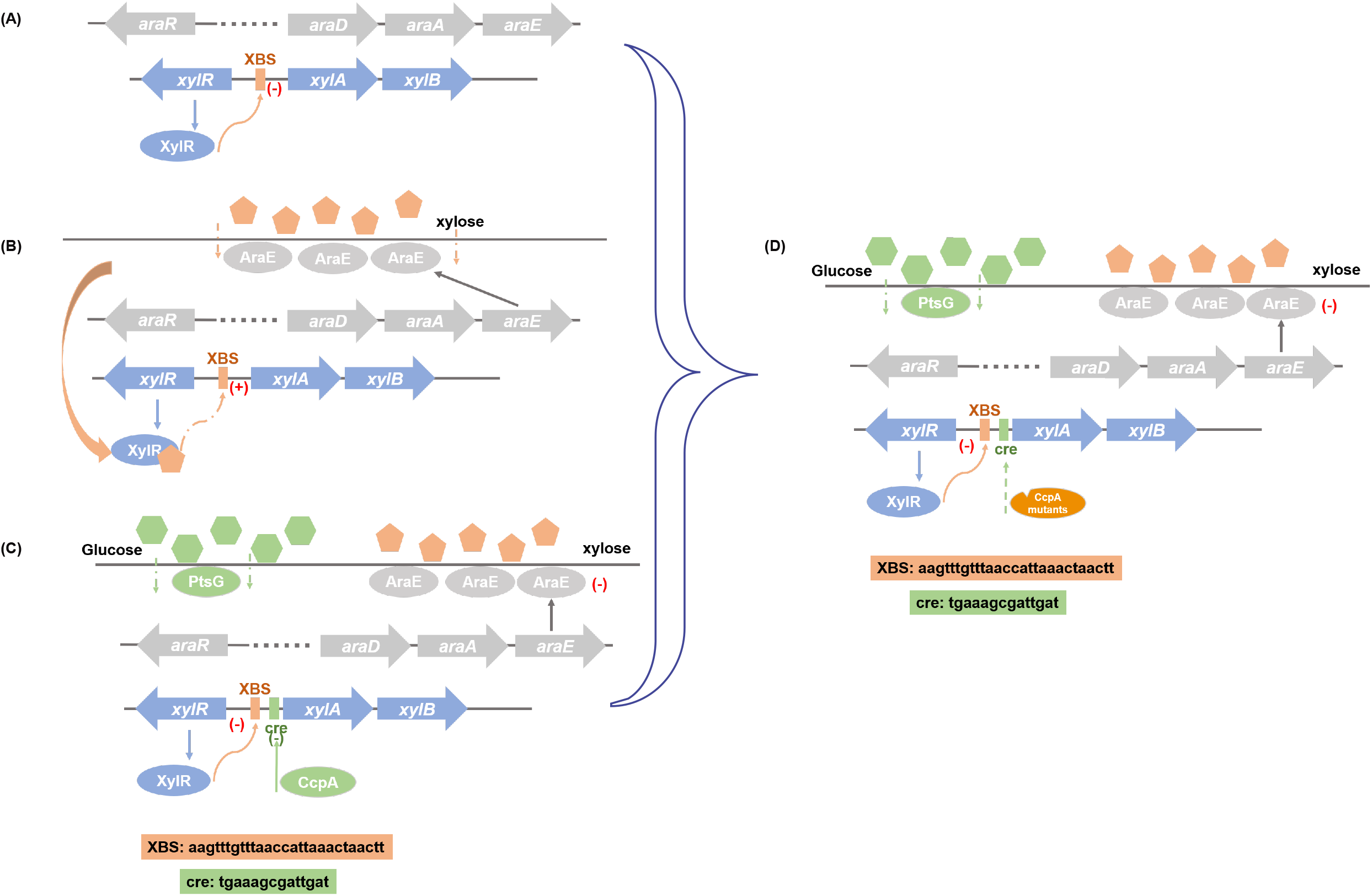
Model of xylose utilization and repression by glucose in *B. licheniformis*. (A) In the absence of xylose, the activity of *XylA* and *XylB* which responsible for xylose utilization were repressed by XylR. (B) In the presence of xylose, the expression of *XylA* and *XylB* were activated. (C) In the exists of xylose and glucose, glucose activates CcpA binds with cre sites. The expression of *XylA* and *XylB* were repressed. (D) In our previous study has demonstrated that the binding ability of CcpA mutants with cre sites have changed. These changes of CcpA maybe affect the xylose utilization at the exist of glucose.

**FIG 6.**
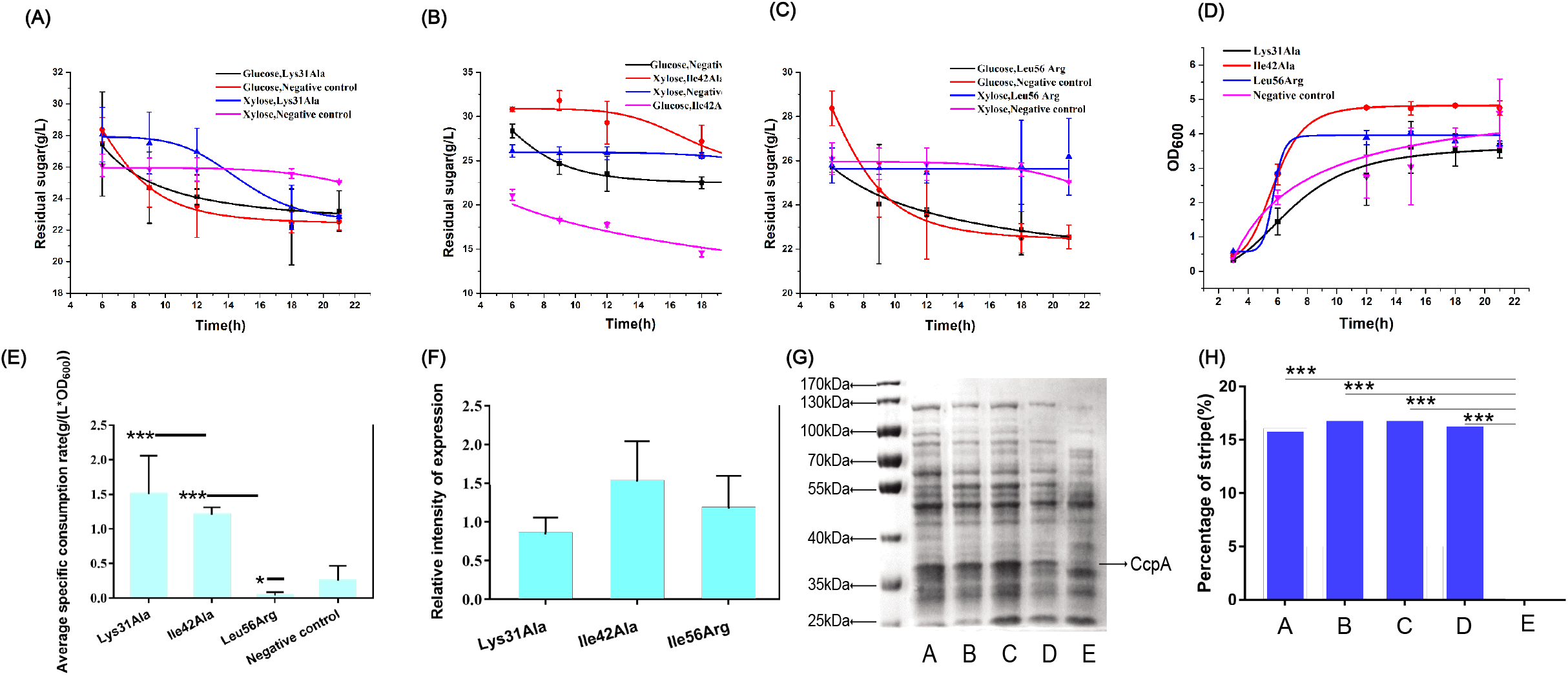
Sugar consumption and biomass of overexpress CcpA mutant strains in fermenting glucose and xylose mixture (approximate 30 g/L glucose and 30 g/L xylose). All experiments were performed in triplicate. (A) (B) (C) Sugar consumption of overexpress CcpA mutation strains in fermenting glucose and xylose mixture. (D) Biomass of overexpress CcpA mutation strains in fermenting glucose and xylose mixture. (E) Xylose average specific consumption rate of overexpress CcpA mutation strains in fermenting glucose and xylose mixture after cultured 21h. The formula for calculating the xylose average specific consumption rate was xylose consumption divided by OD600 (F) The expression level of *ccpA* mutants in in fermenting glucose and xylose mixture. (G) Intracellular SDS-PAGE of B. licheniformis CA, which contain different CcpA mutants. A: Intracellular SDS-PAGE of B. licheniformis CA, which contains Lys 31 Ala. B: Intracellular SDS-PAGE of B. licheniformis CA, which contains Ile 42 Ala. C: Intracellular SDS-PAGE of B. licheniformis CA, which contains Leu 56 Arg. D: Intracellular SDS-PAGE of B. licheniformis CA, which contains origin CcpA. E; Intracellular SDS-PAGE of B. licheniformis CA. (H) The proportion of CcpA mutants intracellular of B. licheniformis CA expressing different mutants. ***——P<0.001, **—P<0.01, *——P<0.05.

To confirm that these differences in xylose and glucose utilization were caused by the mutation of CcpA rather than the expression level, the expression levels of CcpA mutants were determined. Transcriptional strength and the translational levels were detected as is shown in Fig. 6F, Fig. 6G and Fig. 6H. The transcriptional level was detected by qPCR. Compared with wild-type CcpA, the relative values of transcription strength of CcpA mutants did not have significate difference. What is more, the ratio of the CcpA mutants and wild-type CcpA bands were calculated as shown in Fig 6H. This result demonstrates that the mutation of CcpA did not change the CcpA expression level. These findings demonstrate that the Lys31Ala, Ile42Ala, and Ile56Ala CcpA mutants affected the ability of CcpA to bind cre sites, which is responsible for the utilization of xylose.

## Discussion

A large part of coding capacity in the species of *Bacillus* is dedicated to carbohydrate uptake and metabolism, depending on species and functional gene assignment (34). These estimates reside in close association with the well-documented ability of *Bacillus* to utilize a diverse array of carbohydrates (35). The LacI family regulator is representative of a class of carbohydrate metabolism-related transcription factors, and CcpA is the most studied one (29). Approximately 60 residues of the DNA-binding domain are present in its N-terminus, other domains of CcpA are mainly responsible for recognition with cofactors such as HPr and Crh (36). In previous studies, the function of CcpA was affected by the amino acid substitutions, the mutation of Val302 in *Clostridium acetobutylicum* change the priority of xylose and glucose (6). However, these mutants have mainly focused on sites within cofactor binding domain(2, 6, 30) or those highly conserved amino acids within the DNA-binding domain(2, 29). In the current study, we experimentally validated the effect of multiple sites within the DNA-binding domains on regulatory functions of CcpA. It appears that alanine substitution of Lys31 and Ile42, located within the 3^rd^ helices of the DNA-binding domain, lead to alleviated repression of xylose utilization in the presence of glucose. Additionally, the Leu56Arg mutant in the 4^th^ helices exhibited an increased selective utilization of glucose over xylose. These results suggested that changes in microstructure around these domains also contributed to modified regulatory functions of CcpA. This is useful for the engineering of other LacI family regulators.

We found three CcpA mutants with different binding abilities according to the results of rapid in vitro screening guided by homology modeling. According to the available reports, the main reason for xylose utilization repression in the presence of glucose is the binding of CcpA with cre sites and the repressed expression of *XylA* and *XylB* (35). Importantly, the recognition of CcpA with cre sites is structure-dependent. On the one hand, the substructure of CcpA is flexible. The relative distances between different substructures will change during the combination of CcpA between cre sites and cofactors (37). In the current study, when the mutants were expressed in *Bacillus licheniformis* CA, as expected, the rate of xylose utilization is distinctly different. These differences were mainly attributed to the different binding abilities, which was supported by in vitro characterization. The mutation sites selected were distributed in different substructures of the DNA-binding domain, which had implications for the flexibility of the substructure. Previous study had demonstrated that electrostatic interactions between the proteins and the DNA and steric hindrance effect the binding of DNA and protein (38, 39). In our study, that of Lys31Ala and Ile42Ala is small. Therefore, CcpA mutants had a much weaker binding to cre sites. In comparison, the steric hindrance of Leu56Arg was apparently larger. Therefore, Leu56Arg had a higher binding capacity. These results proved that in vitro interaction between the regulator and the target genes could offer credible evidence and be helpful for engineering regulatory proteins.

In the last few years, the function of CcpA has been intensely investigated (40, 41). Many works have been done (42, 43), to explore the relationship of CcpA between structure and function. For example, the amino acids were mutated to explore the activation (*alsS*, *ackA*) and repression (*xynP*, *gntR*) regulated by CcpA in *B. subtilis* (3). The preference for xylose utilization was changed due to the amino acid mutation at positions 302 of CcpA in *Clostridium acetobutylicum* (6). In both studies, amino acids selected were located at the cofactor binding domain. In our study, the preference of xylose was changed because of the amino acid substitutions located in the 3^rd^ and 4^th^ helix in DNA-binding domain, which was a subdomain with few reports. Based on the current results, this substructure might be one of the key domains for recognition and DNA-binding. Furthermore, the “in vitro interaction-in vivo characterization” strategy we used in this study to screen 30 CcpA mutants was quick and cost-effective, compared with in vivo screening alone in previous studies (3, 6). Although these mutants were not sufficient to comprise a mutation library, this method can be easily applied in a large-scale mutagenesis screen.

In conclusion, we have identified three key amino acids, Lys31, Ile42 and Ile56, in the DNA-binding domain of CcpA that affect xylose utilization rate in the presence of glucose in *B. licheniformis*. These changes in xylose utilization have potential uses in fermentation with lignocellulosic biomass. Moreover, because of the functional diversity of CcpA, many metabolic processes were regulated including Biofilm formation and central metabolism (44). These results are helpful for understanding of how microorganisms can flexibly sense and adapt to changes in the external environment. In addition, the strategy used in this study provided a new insight for the engineering of regulatory proteins in *B. licheniformis*, which contributed to the rational design of this important industrial microorganisms.

## Materials and methods

### Bacterial strains and culture conditions

Table 1 lists the bacterial strains and plasmids that were used or generated during this study.

All experiments were performed with three replicates. *B*. *licheniformis* CA is a gene knockout expression host lacking the gene expressing the global regulatory protein CcpA. This strain was designed to explore the influence of CcpA and CcpA mutants. The osmotic medium and agents used for protoplast transformation were prepared according to Waschkau et al and included SMMP medium, SSM buffer, no.416 medium, and sugar (45). *E. coli* were grown at 37 °C at 200 rpm in Luria-Bertani (LB) medium containing 10 g/L tryptone, 5 g/L yeast extract and 10 g/L NaCl. *Bacillus* was grown at 37°C and 250 rpm in LB. TB medium containing 30 g/L xylose and 30 g/L glucose was prepared and used to evaluate the effect of CcpA protein mutants on xylose consumption. Ampicillin (100 μg/mL), 30 μg/mL kanamycin or 20 μg/mL tetracycline was added to the medium when *E. coli* and *Bacillus* were cultivated. The fermentation medium was divided into 250 mL shake flasks, with each flask containing 30 mL of medium, and shaken at 250 rpm. Samples were taken every three hours to determine the OD600 and glucose and xylose content.

### Homology modeling of *B. licheniformis* CcpA and homology comparison

To explore the effect of CcpA on xylose utilization in *B. licheniformis*, a homology model of CcpA from *B. licheniformis* was generated with the SWISS-MODEL server (46). The amino acid sequence of CcpA, which was downloaded from NCBI, was the server input. The amino acids of CcpA from *B. licheniformis* showed high identity (78.55%) with the model in the server. The homology of CcpA from *B. licheniformis* and other closely related bacilli was compared with DNAMAN. All the sequences were downloaded from NCBI.

### CcpA fixed point mutants

The mutated sites and primers used are shown in the Table 2. PCR site-directed mutagenesis was used for mutants (47). Each PCR sample contained 50 nM of each primer (Sangon Biotech, Shanghai, China), 50 mL of Phanta Max DNA polymerase (Vazyme Biotech, Nanjing, China), and 50 ng of template DNA (recombinant plasmid pET28aPA). All the PCR products were purified and digested with *DpnI*, followed by transformation into *BL21(DE3)*. The positive bacteria were cultured in LB medium containing 30 μg/mL kanamycin overnight at 37 °C. Then, the plasmids were extracted and sequenced, and the correct mutant was selected.

### Expression and purification of mutant proteins

Recombinant *BL21(DE3)* bacteria were cultivated in a 250-mL flask containing 30 mL of fermentation medium. First, the strains were grown overnight at 37°C while shaking in LB medium, after which 3% of the medium volume was added to the fermentation medium. IPTG was added to a final concentration of 0.1 mM as an inducer when the OD_600_ of the culture had reached approximately 0.6. Then, the shake flask was transferred to 30°C based on the methods of Min’s (48). After 12 h, the bacteria were collected and ultrasonically broken. The target protein was purified with a Mag-Beads His-tag protein purification kit (Sangon Biotech, Shanghai, China). The purified protein was detected by sodium dodecyl sulfate-polyacrylamide gel electrophoresis (SDS-PAGE) (10% gel) according to the method of Li’s (33).

### Screening of mutant proteins

*B. licheniformis* CcpA and mutant CcpA proteins were expressed and purified as described in our previous studies. Fluorescence polarization immunoassay (FPIA) and electrophoretic mobility shift assay (EMSA) were used to screen and identify the ability of CcpA mutant proteins to bind cre sites. The 5’ terminal fluorescently (Alexa Fluor 488) labeled probe (The sequence was shown in the supplementary information) dsDNA used in the FPIA experiment was obtained by PCR using the *B. licheniformis* genome as a template and the primers CreF/CreR, which are shown in Table 2. Before testing with a multifunctional enzyme marker (BioTek Instruments, Winooski, VT), 100 nM fluorescently labeled probe dsDNA was incubated with 60 μg of mutant CcpA protein in binding buffer (60μL) consisting of 25 mM Tris-HCl, 3 mM NaCl, 3 mM MgCl_2_ and 0.1 mM DTT according to the method of Xu, H Q et al at room temperature for 20 min (49). Then, the total volume was added to 100 μL by buffer, and we measured the excitation at 485 nm and the absorption at 528 nm with black 96-well plates with multi-detection microplate reader (BioTek, USA). The DNA probes used in the EMSA were amplified by PCR with CreF1/CreR using the *B. licheniformis* genome as a template, and the 3’ DNA probe was labeled with biotin. Before electrophoresis, 30 ng of DNA probe was incubated with 3 μg of mutant CcpA protein in binding buffer (chemiluminescent EMSA kit, GS009, Beyotime, Shanghai, China) at room temperature. All the next steps were conducted as indicated in the kit instructions. Bio-Rad Mini-Protean III electrophoresis apparatus and Bio-Red Mini Trans-Blot (Bio-Rad, California, USA) were use in this study.

### *CcpA* deletion and CcpA mutation overexpression

To eliminate the effect on the evaluation of CcpA mutation, the original genetic of *ccpA* was deleted. The bacteria and plasmids are shown in Table 1. The medium and reagents for *B. licheniformis* and *E. coli* were based on Li’s methods (50). *E. coli* and *B. licheniformis* were cultured in LB medium. The recombination plasmids TC and TCFKFC were used to construct gene knockout cassettes. Then, the gene knockout cassette was enzyme digested by *Kpn*I and *Xho*I. Then, the knockout cassette was linked with plasmid pT. Last, the recombination plasmid pTCFKFC was transformed into *B. licheniformis* according to Li’s methods (33). The next steps were performed according to Wang’s methods (51). The original genetic mechanism of *ccpA* was inactivation because of the insertion of foreign genes (*kanamycin resistance gene* (MK784777.1)).

The *ccpA* overexpression plasmids were constructed based on the pHY300-PLK vector using the primers listed in Table 2. All molecular experiments were performed according to standard molecular cloning protocols. To construct recombinant plasmids for the expression of CcpA and mutant CcpA proteins, we first amplified the P43 promoter using the Taq enzyme (Vazyme Biotech, Nanjing, China) with the primers T-p43-F/T-p43-R listed in Table 2 and then ligated the amplification product to pMD18-T. The target genes *ccpA*-Lys-31-Ala, *ccpA*-Ile-42-Ala and *ccpA*-Lue-56-Arg were acquired by PCR with the primers T-BL2-CcpA-F/T-BL2-CcpA-R, which are listed in Table 2, and the template, which is listed in Table 1. The target gene *ccpA* was acquired by PCR with the primers T-BL2-CcpA-F/T-BL2-CcpA-R, and the template was the *B. licheniformis* genome. The primer pair T-P43-ReF/T-P43-ReR was used to linearize the TP plasmid. Then, the target fragment and TP fragment were assembled with a ClonExpress Ultra One Step Cloning Kit (Vazyme Biotech, Nanjing, China). Fragments for P43, CcpA and the CcpA mutants were amplified, purified and digested with *Bgl*II and *Eco*RI, followed by incorporation into pHY300-PLK. Then, the recombinant plasmids were individually transformed into *ccpA*-defective *B. licheniformis* strains by electrotransformation and cultured overnight at 37°C in solid medium containing 20 μg/mL tetracycline. The plasmids were extracted and sequenced to ensure the correct transformants.

### Carbon source consumption measurements

Strains harboring different mutants were individually inoculated into LB medium containing tetracycline and cultured at 37 °C, 250 rpm overnight. Then, the cultures were inoculated at OD_600_ = 0.15 into fresh 30 mL TB medium supplemented with glucose (30 g/L) and xylose (30 g/L). Samples of the CcpA-defective strains overexpressing the CcpA mutant were collected at different time points (6 h, 9 h, 12 h, 15 h, 18 h and 21 h) for assessment. One milliliter of each sample was centrifuged at 12000 rpm for 10 min, and an equal volume of 10% trichloroacetic acid was added to the supernatant to remove impurities. Carbon source consumption were assessed by HPLC (Thermo Fisher Scientific, Shanghai, China) with a Polyamino HILIC (Dikma, Beijing, China) chromatographic column according to the manufacturer’s specifications (52, 53).

### The expression level of *CcpA* mutants

To verify that the CcpA mutants did not change the expression level of *ccpA* and thus affect the utilization of xylose. Strains harboring different mutants were individually inoculated into LB medium containing tetracycline and cultured at 37 °C, 250 rpm overnight. Then, the cultures were inoculated at OD600 = 0.15 into fresh 30 mL TB medium. Strains harboring *ccpA* was the negative control. The expression levels of the *ccpA* mutants were determined by real-time quantitative PCR (RT-qPCR). Samples were collected at 8 h, and the cells were harvested after isolation. Total RNA was extracted with the Simply Ptotal RNA extraction kit (Bioflux, Beijing, China) according to the instructions and quantified with a Quawell Q5000 ultraviolet-visible spectrophotometer (Quawell Technology, San Jose, CA). cDNA was prepared according to the instructions of the PrimeScript RT reagent kit (Vazyme Biotech, Nanjing, China) and used as a template for analysis by real-time PCR with a real-time PCR system with ChamQ™ Universal SYBR qPCR Master Mix (Vazyme Biotech, Nanjing, China) and the primers q*ccpA*-F/q*ccpA*-R. The internal reference gene was *rpsE* (20). The relative transcript strength was calculated using the 2^−ΔΔCt^ method (33). Translation levels of CcpA mutants were measured by the ratio of the CcpA mutant bands. *B. licheniformis* harbored CcpA mutants were cultured with TB medium suppled 30 g/L glucose and 30 g/L xylose. The strains were collected after 24 hours. The strains were washed twice with PB buffer, and then resuspended in PB buffer containing 10g/L lysozyme. The bacterial solutions were incubated for 1 hour at 37 °C and broken by sonication on ice and then centrifuged at 12000 rpm for 10 min. Twenty μg of total protein per sample was loaded onto a gel.

## Funding

This work was supported by National Key Research & Development Program of China (2020YFA0907700, 2018YFA0900300 and 2018YFA0900504), the National Natural Foundation of China (31401674), the National First-Class Discipline Program of Light Industry Technology and Engineering (LITE2018-22), and the Top-notch Academic Programs Project of Jiangsu Higher Education Institutions.

---

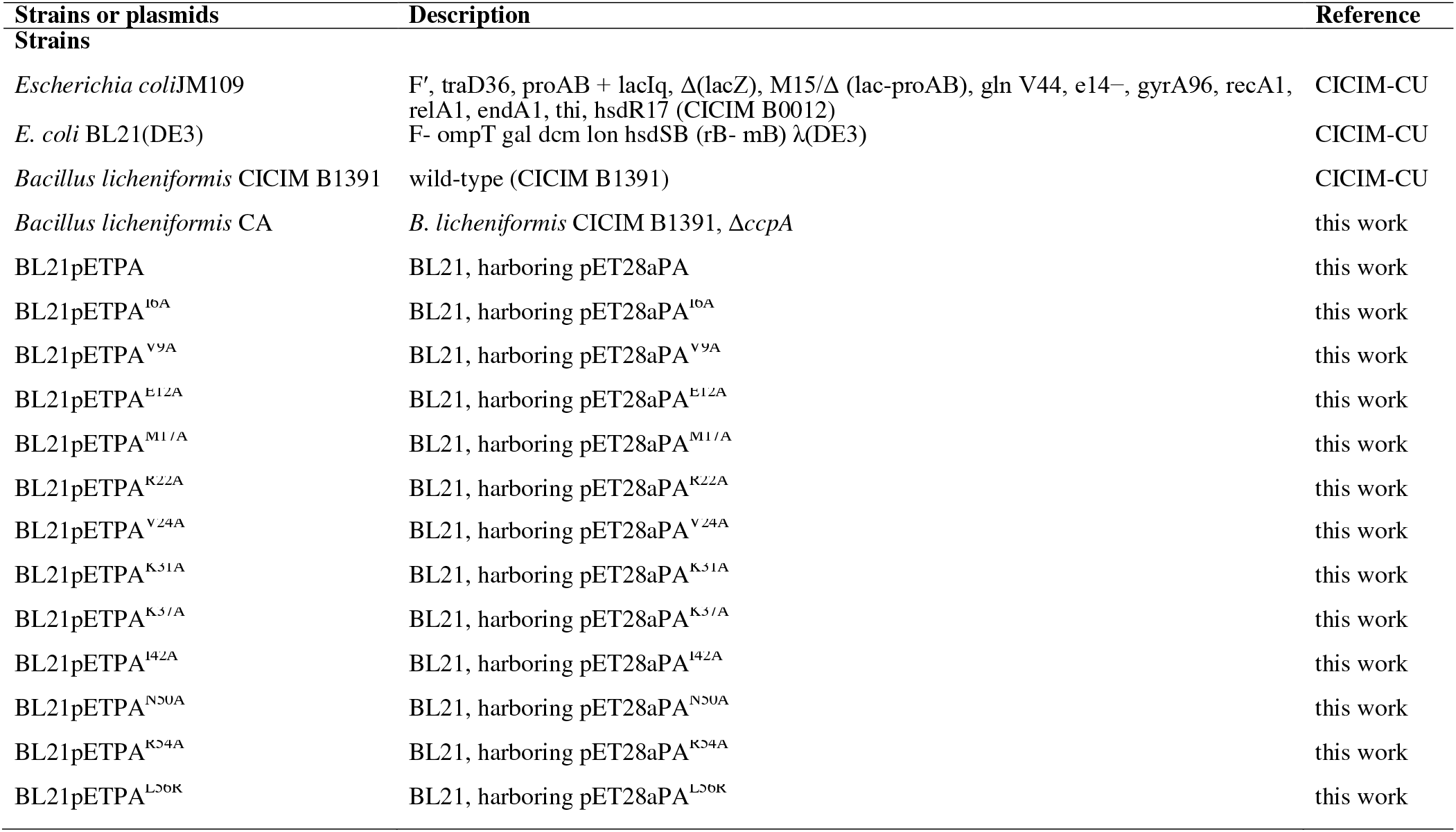
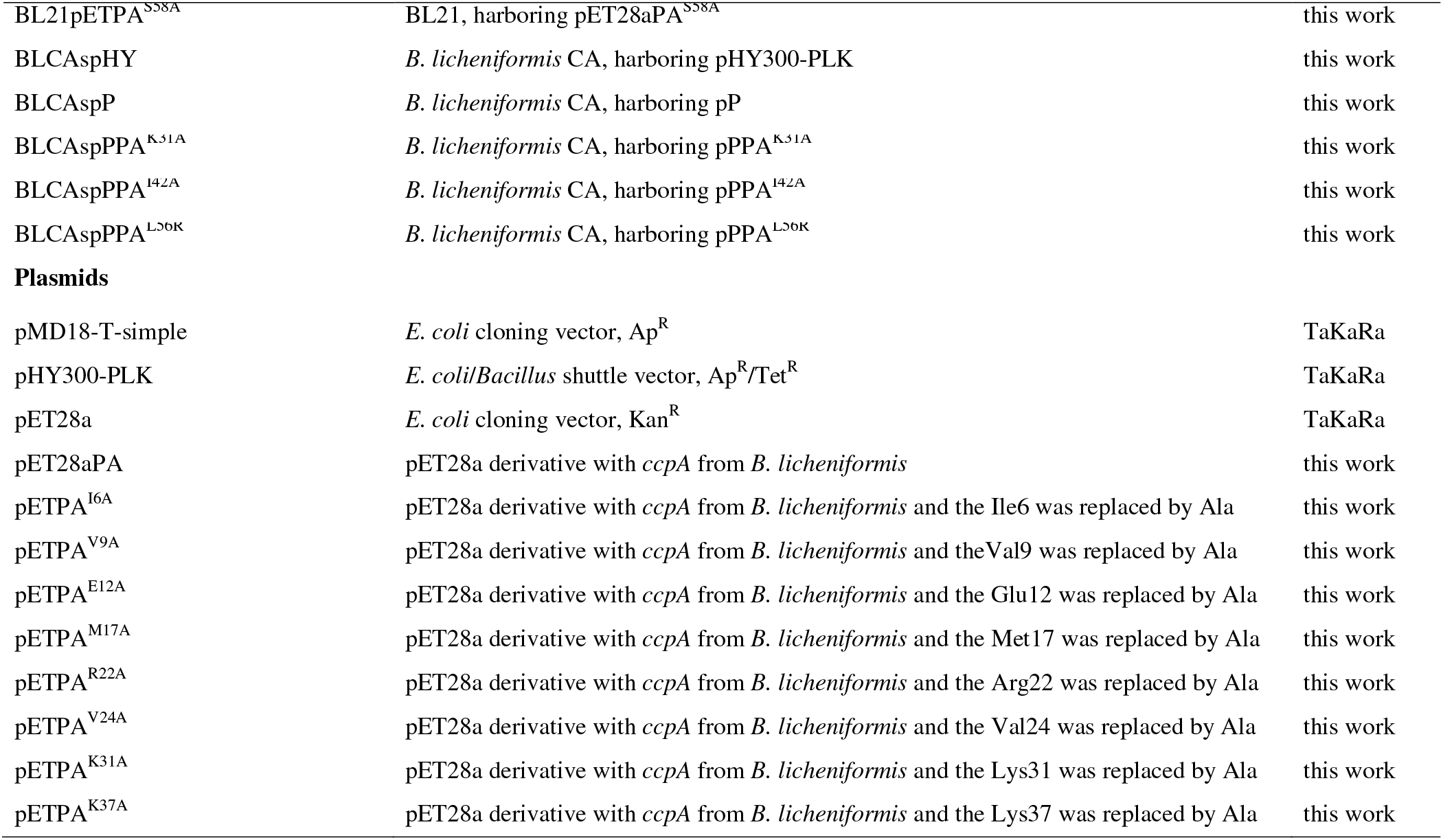
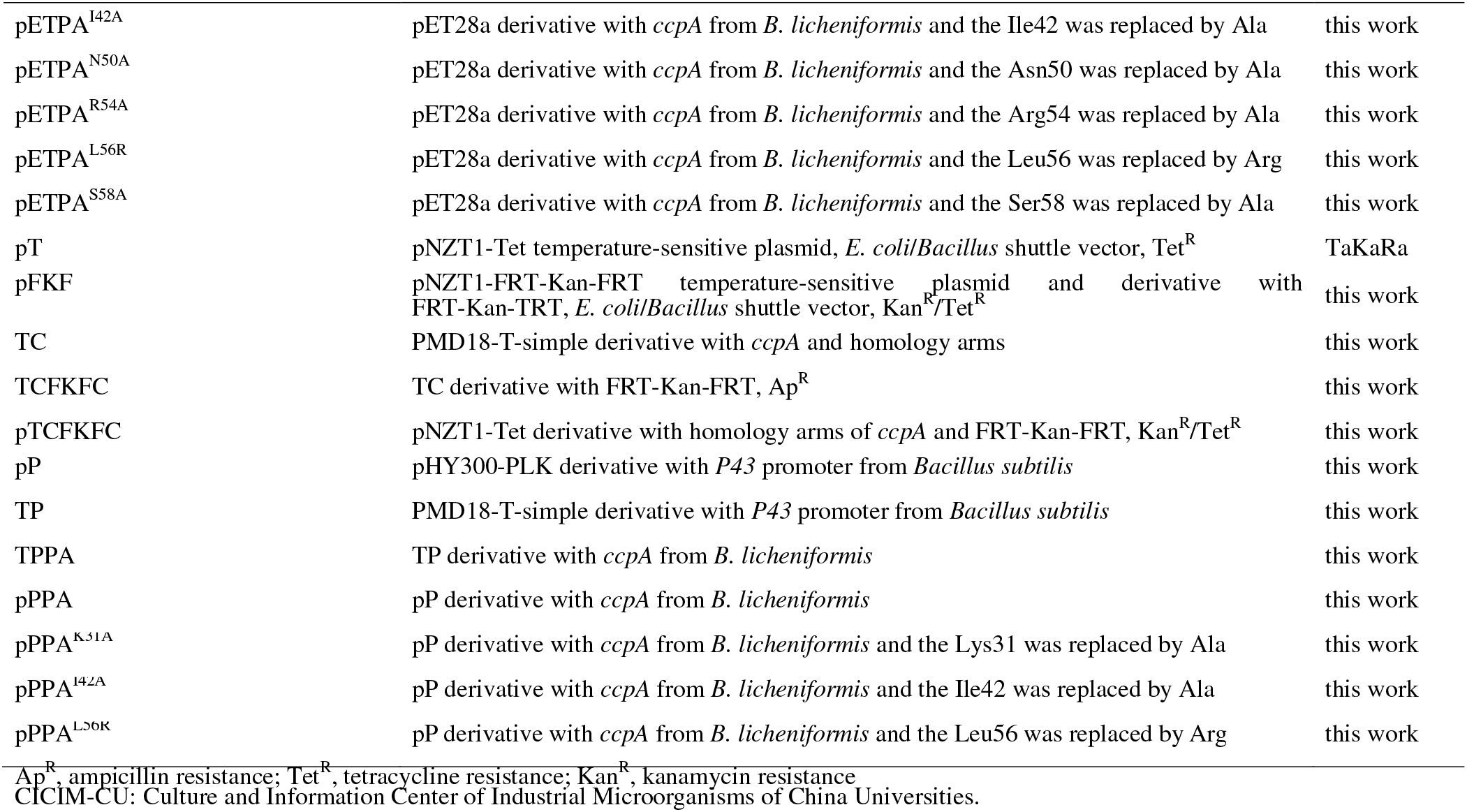

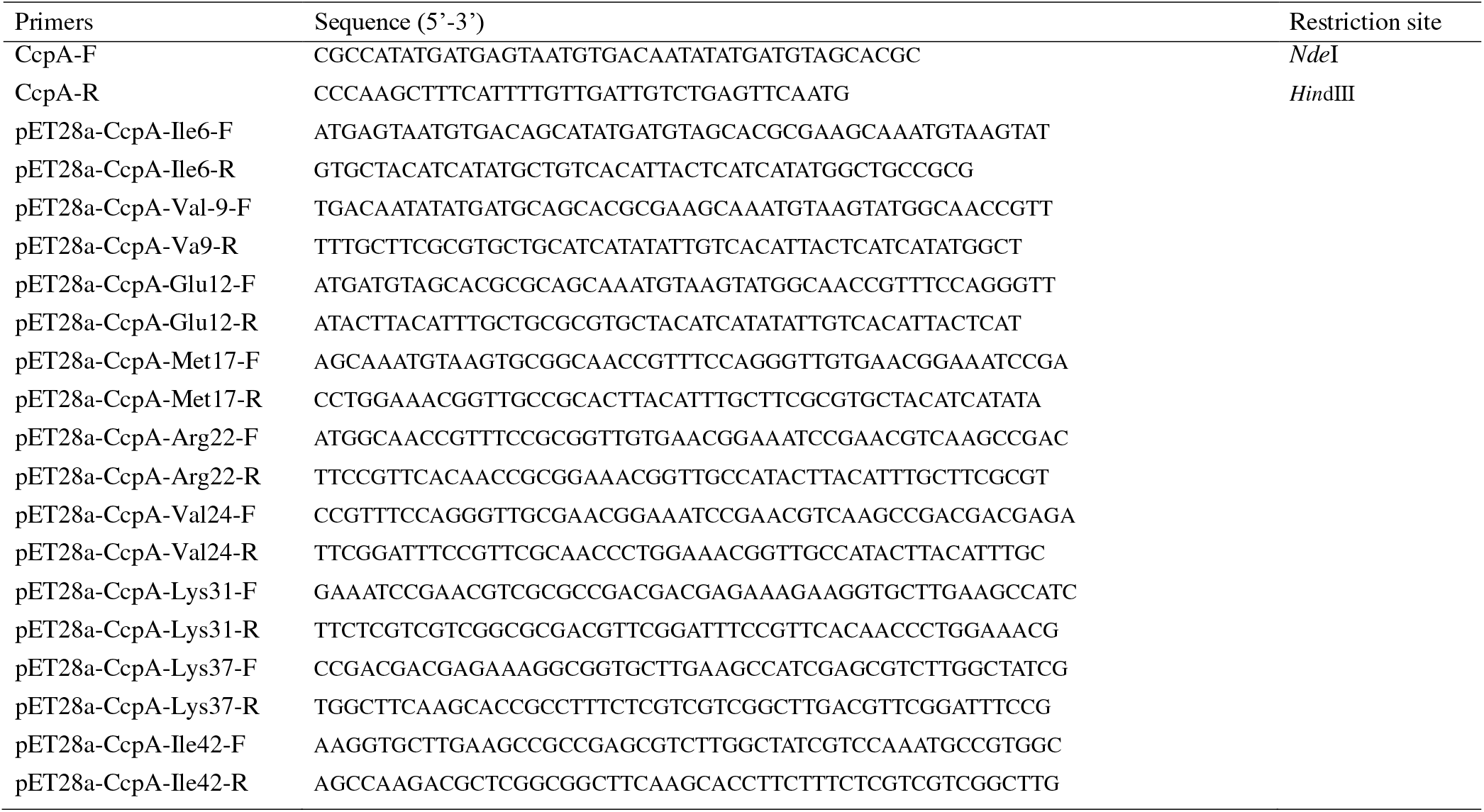
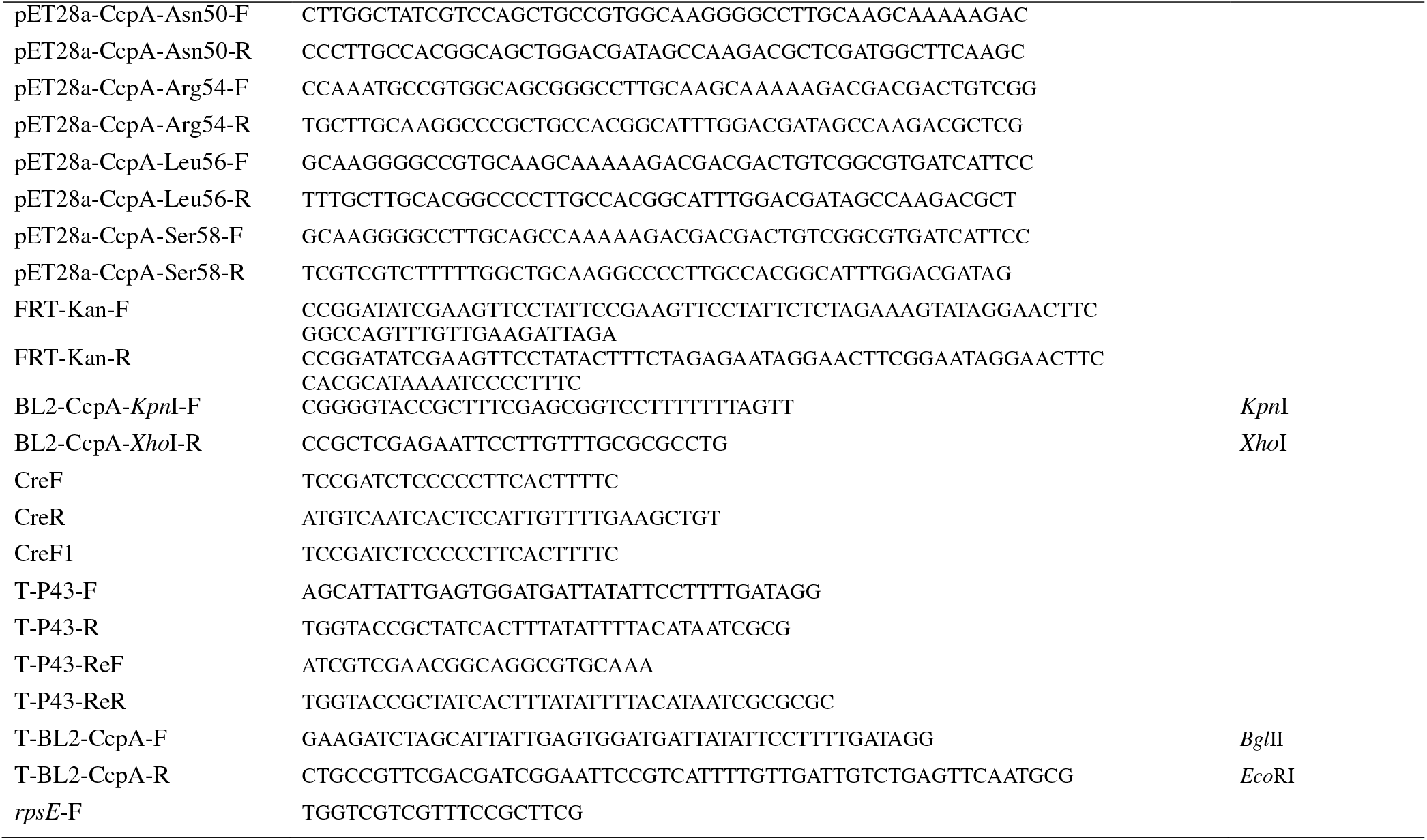
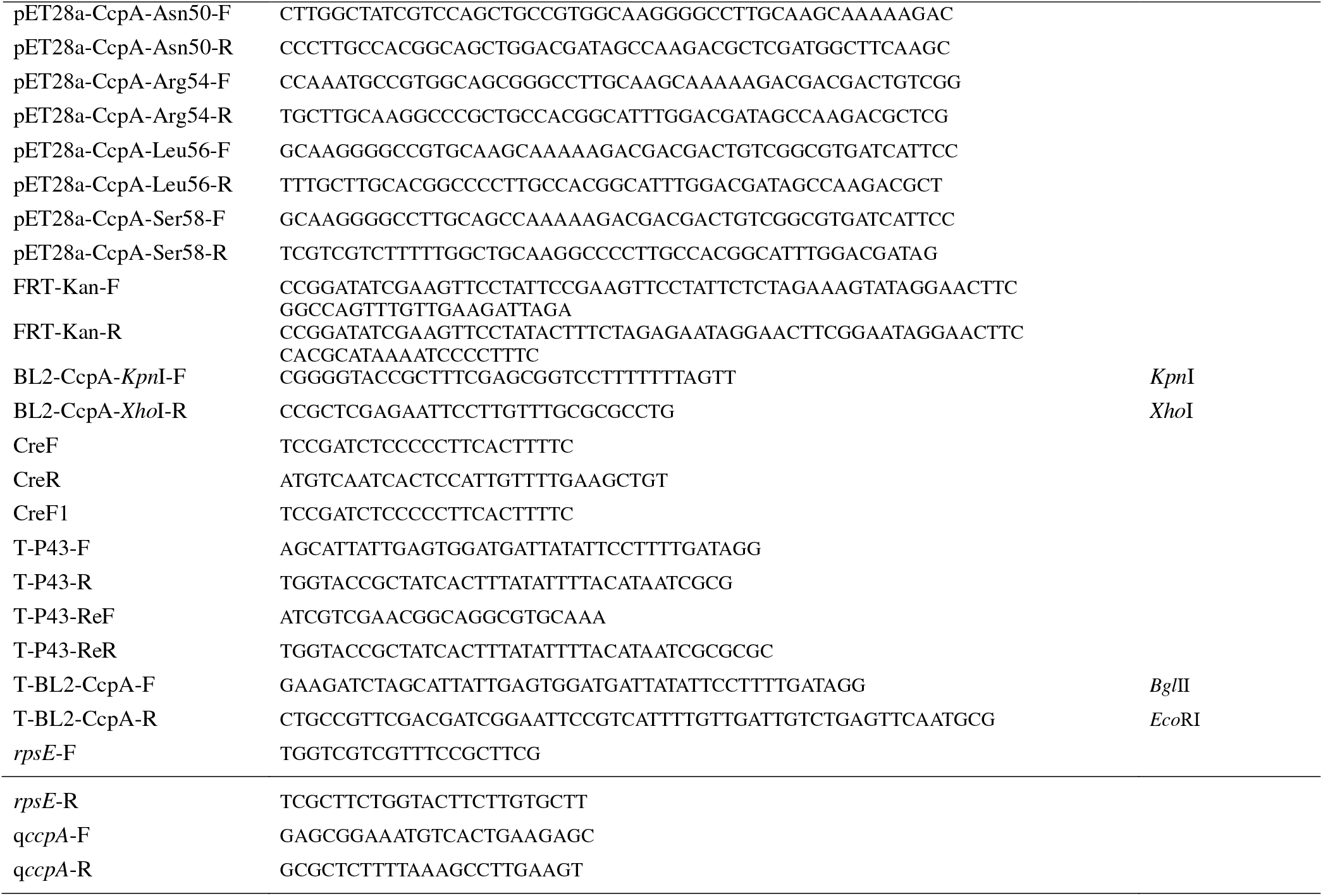
Primers Used to Construct Recombinant Plasmids

